## Supplementary Methods, Figures and Tables for "Single cell transcriptome analysis of cavernous tissues reveals the key roles of pericytes in diabetic erectile dysfunction"

**One-sentence summaries**: LBH, a novel marker of cavernous pericytes, influences angiogenesis and neurovascular degeneration, thereby preventing diabetic erectile dysfunction.

Seo-Gyeong Bae^1†^**,** Guo Nan Yin^2†^, Jiyeon Ock^2†^, Jun-Kyu Suh^2*^, Ji-Kan Ryu^2,3*^, Jihwan Park^1*^

^1^School of Life Sciences, Gwangju Institute of Science and Technology (GIST), Gwangju, Korea

^2^National Research Center for Sexual Medicine and Department of Urology, Inha University School of Medicine, Incheon, Korea

^3^Program in Biomedical Science & Engineering, Inha University, Incheon, Korea

†These authors contributed equally to this work.

*Corresponding authors

**Jun-Kyu Suh, MD, PhD**

National Research Center for Sexual Medicine and Department of Urology, Inha University School of Medicine

100 Inha-ro, Michuhol-gu, Incheon, Republic of Korea, 22212

**Ji-Kan Ryu, MD, PhD**

National Research Center for Sexual Medicine and Department of Urology, Inha University School of Medicine

100 Inha-ro, Michuhol-gu, Incheon, Republic of Korea, 22212

**Jihwan Park, PhD**

School of Life Sciences, Gwangju Institute of Science and Technology

123 Cheomdangwagi-ro, Buk-gu, Gwangju, Republic of Korea, 61005

**List of Supplementary Materials**

Supplementary Materials and Methods

Fig. S1. Chondrocyte subsets in single cell RNA sequencing data of mouse cavernous tissues.

Fig. S2. Volcano plots showing DEGs between diabetic and normal conditions in each cell type.

Fig. S3. Gene ontology high in normal compared to diabetes in each cell type.

Fig. S4. Gene ontology high in diabetes compared to normal in each cell type.

Fig. S5. Pericyte-specifically expressed genes in single-cell RNA sequencing data.

Fig. S6. Expressions of *Lbh* in single-cell RNA sequencing data from tissues other than the penis.

Fig. S7. Cell-cell interactions between pericytes and other cell types showing significant differences in between normal and diabetes.

Fig. S8. Cell-cell communication between cell types in normal and diabetes using CellChat.

Fig. S9. RT-PCR validation of differentially expressed genes from single cell RNA sequencing analyses.

Fig. S10. LBH immunofluorescence staining in corpus cavernosum tissues after infection with lentiviruses containing ORF mouse clone of *Lbh*

Fig. S11. LBH enhances pericyte angiogenesis and MPG neurite sprouting under high glucose conditions.

Fig. S12. Expression of marker genes of pericyte and smooth muscle cells in human penis single cell RNA sequencing data.

Table S1. Relative density of mouse angiogenesis array spots

Table S2. Physiologic and metabolic parameters: 2 weeks after treatment with PBS, NC, LBH O/E

References

**Supplementary Material and Methods**

**TUNEL Assay**

The MCPs cell death after infected with lentiviruses ORF control particles (NC, 5x10^4^ infection units per millilitre cultured medium; Origene Technology) or ORF clone of mouse *Lbh* (LBH O/E, 5x10^4^ infection units per millilitre cultured medium; Origen Technology) under NG or HG conditions were evaluated by TUNEL (terminal deoxynucleotidyl transferase-mediated deoxyuridine triphosphate nick-end labeling) assay using the ApopTag® Fluorescein In Situ Apoptosis Detection Kit (S7160, Chemicon, Temecula, CA, USA) according to the manufacturer’s instructions and as described previously (Yin et al., 2022). The numbers of TUNEL-positive apoptotic cells were obtained by a confocal fluorescence microscope.

**Cell migration assay**

MCPs migration assays were performed using the SPLScar^TM^Block system (SPL life sciences, Pocheon-si, Gyeonggi-do, Korea) on 60-mm culture dishes as described previously (Yin et al., 2022). In brief, MCPs were infected with lentiviruses ORF control particles (NC, 5x10^4^ infection units per millilitre cultured medium; Origene Technology) or ORF clone of mouse *Lbh* (LBH O/E, 5x10^4^ infection units per millilitre cultured medium; Origen Technology) under NG or HG conditions. Then conditioned cells were seeded into 3-well blocks at >90% confluence. Blocks were removed after 5 hours and cells were incubated for additional 24 hours in DMEM medium containing 2% FBS and thymidine (2 mM, Sigma-Aldrich). The images were taken using a phase-contrast microscope (Olympus), and cell migration was analyzed by determining the percentage of cells that moved into the frame line showed in the figures from four separate block systems in a blinded manner using Image J software (National Institutes of Health [NIH] 1.34, http://rsbweb.nih.gov/ij/).

***In vitro* tube formation assay**

Tube formation assay was performed as described previously (Yin, 2022). MCPs were infected with lentiviruses ORF control particles (NC, 5x10^5^ infection units per millilitre cultured medium; Origene Technology) or ORF clone of mouse Lbh (Lbh O/E, 5x10^5^ infection units per millilitre cultured medium; Origen Technology) under high glucose conditions for at least 3 days. Tube formation assays were performed in a 48-well plate with 100 µL of growth factor-reduced Matrigel (Becton Dickinson). The assay was performed in a CO_2_ incubator and the images were obtained at 24 hours at a screen magnification of 40 with a phase-contrast microscope (CKX41, Olympus, Japan). The numbers of master junctions from four separate experiments was determined by using Image J software.

***Ex vivo* neurite sprouting assay**

MPG tissues were harvested and maintained as described previously (Ghatak et al., 2022). The tissues were covered with matrigel and incubated at 37 °C for 10 minutes in a 5% CO_2_ atmosphere, followed by incubation in 1.2 ml of complete Neurobasal medium (Gibco) containing 0.5 nM GlutaMAX™-I (Gibco) and 2% serum-free B-27 (Gibco). The MPG tissues were infected with lentiviruses ORF control particles (NC, 5x10^4^ infection units per millilitre cultured medium; Origene Technology) or ORF clone of mouse *Lbh* (LBH O/E, 5x10^4^ infection units per millilitre cultured medium; Origen Technology) under NG or HG conditions. Five days later, neurite outgrowth segments were then fixed in 4% paraformaldehyde for at least 30 minutes and immunofluorescence staining was performed with an anti-neurofilament antibody (N4142; Sigma-Aldrich; 1:50). Images were obtained with a phase-contrast microscope (CKX41, Olympus, Japan). Quantitative analysis of neurite length was determined by using Image J software.

**Supplementary Figures**

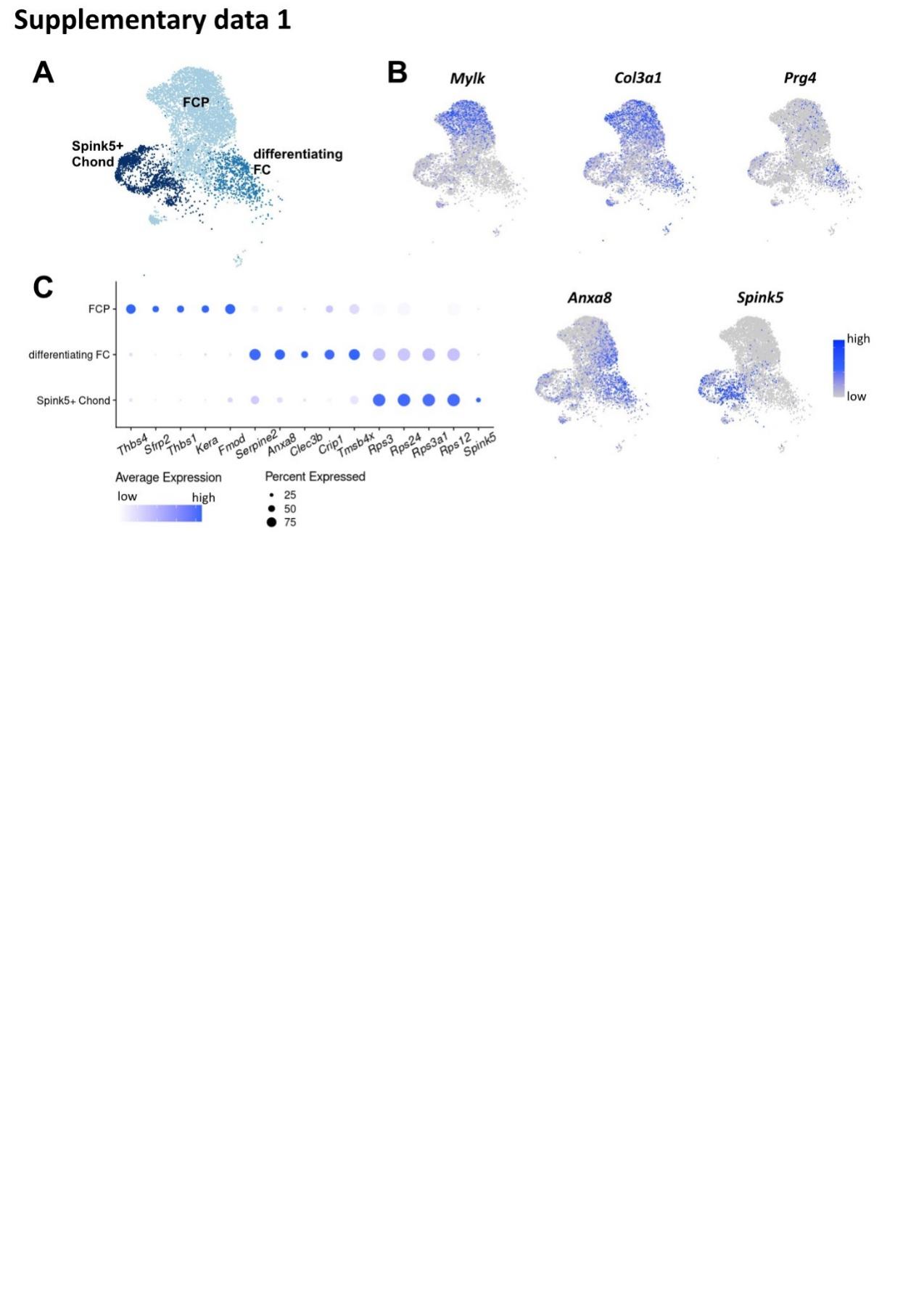

**Fig. S1. Chondrocyte subsets in single cell RNA sequencing data of mouse cavernous tissues.** (**A**) UMAP projection of three chondrocyte clusters in single cell data of mouse cavernous tissues. (**B**) Expression of marker genes of three chondrocyte subsets. *Mylk* and *Col3a1* for FCP; *Prg4* and *Anxa8* for differentiating FC; *Spink5* for *Spink5*+ Chond. (**C**) Dot plot showing the expression of top five marker genes of each chondrocyte subset. The color represents the average expression level, and the size of the dots represents the percentage of cells expressing each gene.

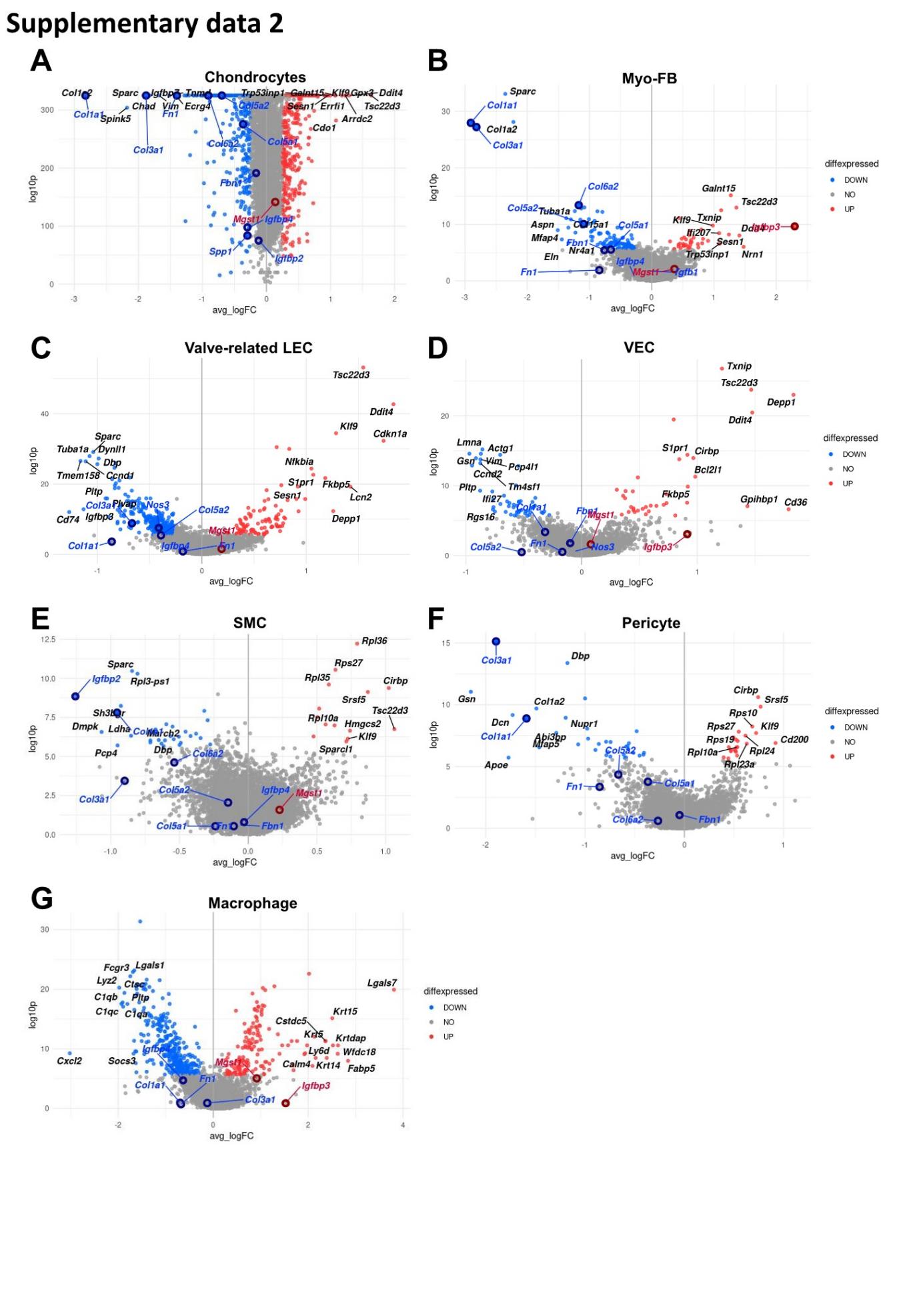

**Fig. S2. Volcano plots showing DEGs between diabetic and normal conditions in each cell typ**e**.** (**A**) DEGs between diabetic and normal conditions in Chondrocytes. (**B**) DEGs between diabetic and normal conditions in Myo-FB. (**C**) DEGs between diabetic and normal conditions in Valve-related LEC. (**D**) DEGs between diabetic and normal conditions in VEC. (**E**) DEGs between diabetic and normal conditions in SMC. (**F**) DEGs between diabetic and normal conditions in Pericyte. (**G**) DEGs between diabetic and normal conditions in Macrophage. The top 10 (based on log-fold change) DEGs are indicated with gene names, and genes identified as having high or low expression in diabetes in previous studies are indicated with gene names in red or blue. DEGs with adjusted p-value > 0.05 were indicated in gray. The LEC was not shown as a volcano plot because there were only two significant DEGs, and Schwann cells have no significant DEG (adjusted p-value < 0.05).

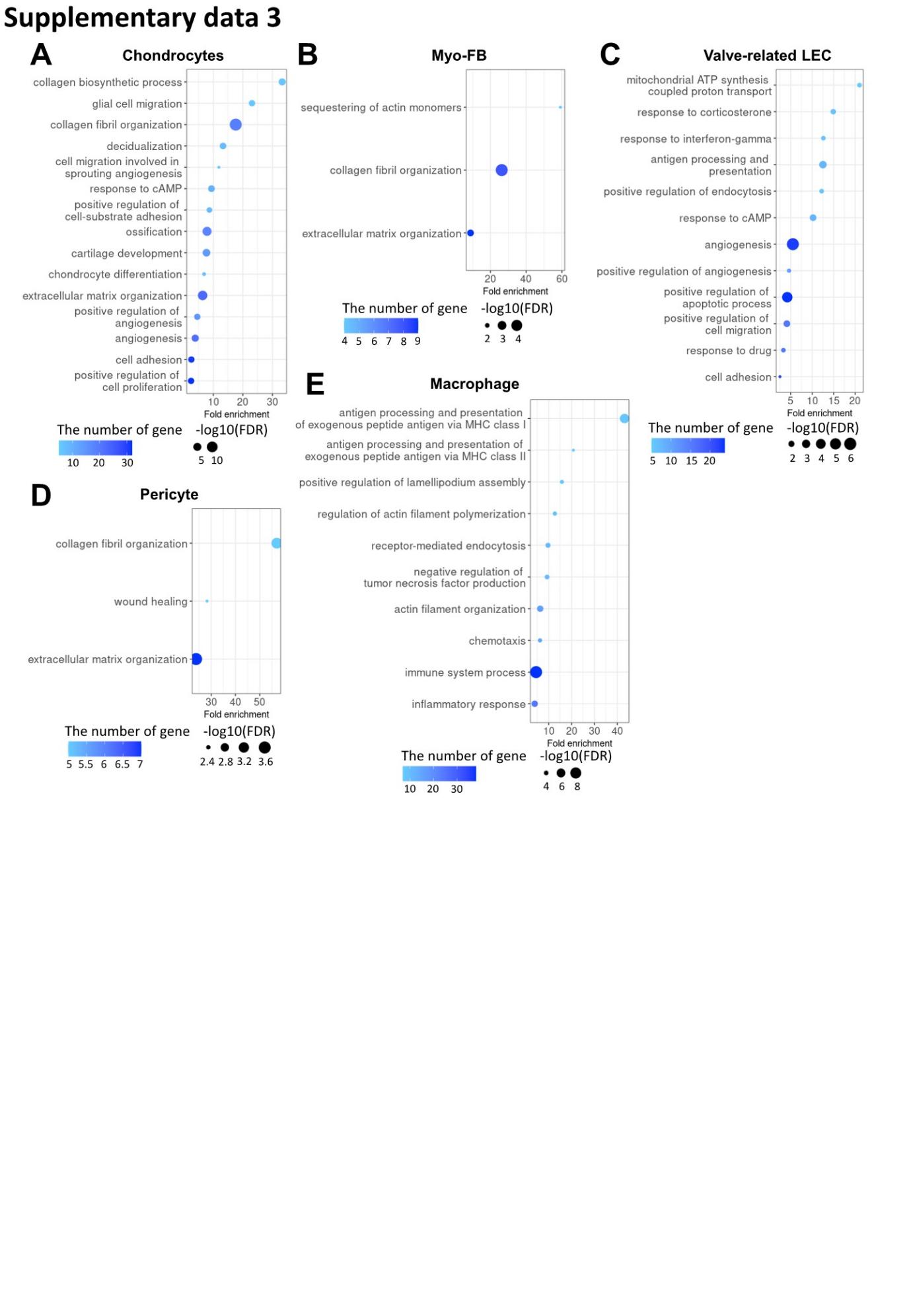

**Fig. S3. Gene ontology high in normal compared to diabetes in each cell type.** (**A**) Gene ontology analysis of the genes higher in normal compared to diabetes in Chondrocytes. (**B**) Gene ontology analysis of the genes higher in normal compared to diabetes in Myo-FB. (**C**) Gene ontology analysis of the genes higher in normal compared to diabetes in Valve-related LEC. (**D**) Gene ontology analysis of the genes higher in normal compared to diabetes in Pericyte. (**E**) Gene ontology analysis of the genes higher in normal compared to diabetes in Macrophage. There was no significant term in LEC, VEC, SMC, and Schwann cells (p-value < 0.05 and FDR < 0.25).

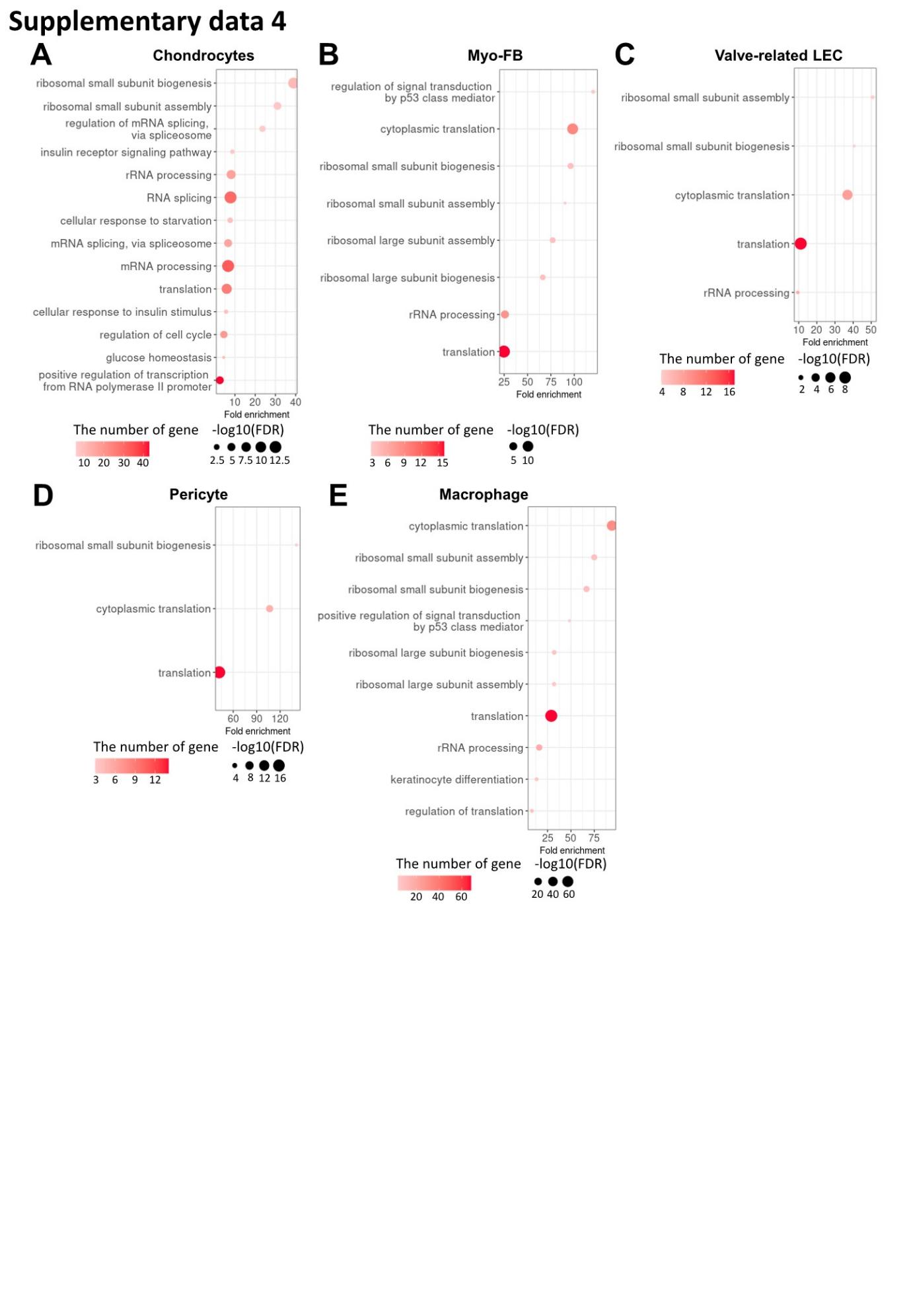

**Fig. S4. Gene ontology high in diabetes compared to normal in each cell type.** (**A**) Gene ontology analysis of the genes higher in diabetes compared to normal in Chondrocytes. (**B**) Gene ontology analysis of the genes higher in diabetes compared to normal in Myo-FB. (**C**) Gene ontology analysis of the genes higher in diabetes compared to normal in Valve-related LEC. (**D**) Gene ontology analysis of the genes higher in diabetes compared to normal in Pericyte. (**E**) Gene ontology analysis of the genes higher in diabetes compared to normal in Macrophage. There were no significant terms in LEC and Schwann cells (P-value < 0.05 and FDR < 0.25), and only translation-related terms were identified in VEC and SMC.

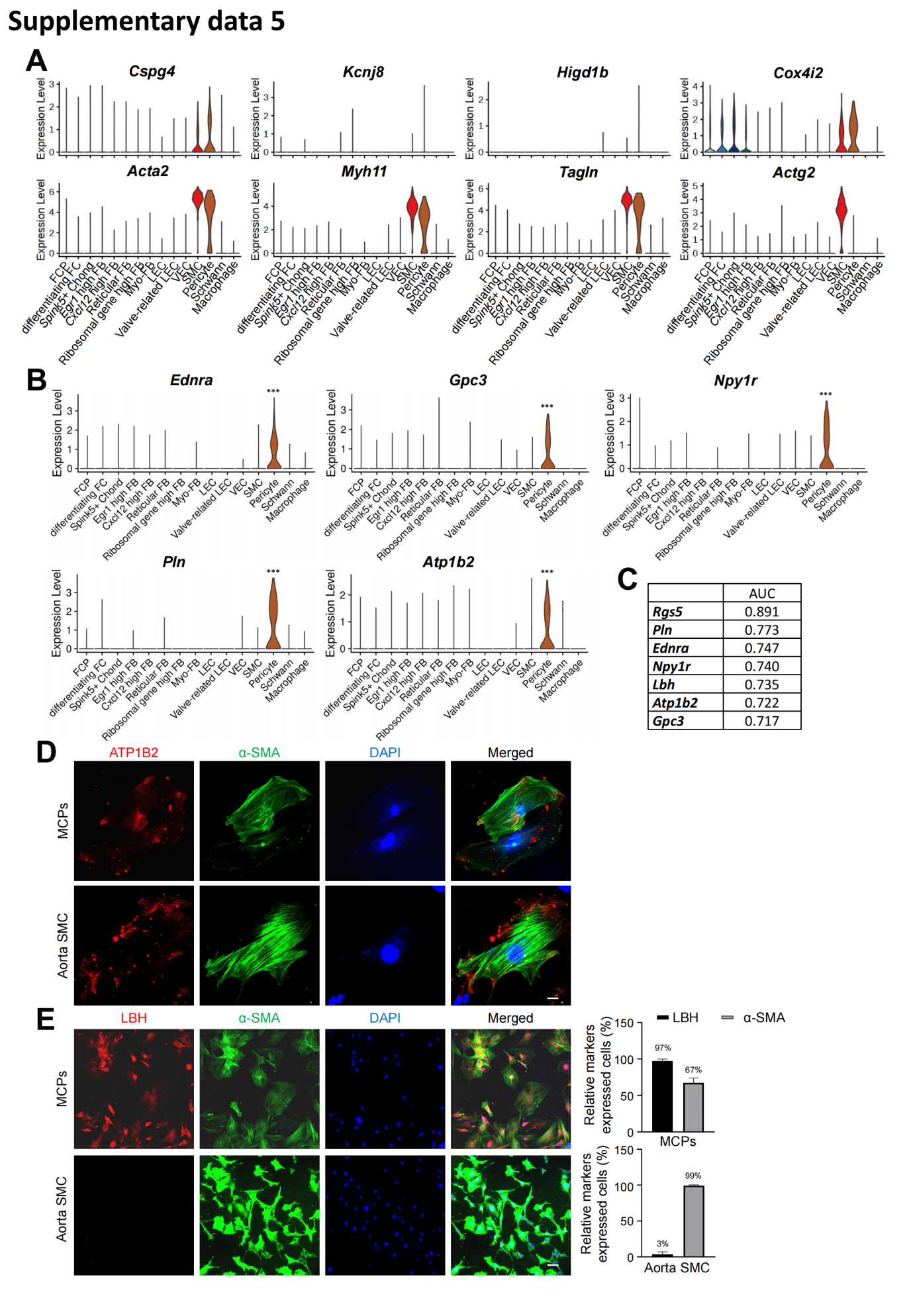

**Fig. S5. Pericyte-specifically expressed genes in single-cell RNA sequencing data.** **(A)** Violin plots showing the expression of known marker genes of pericyte (*Cspg4, Kcnj8, Higd1b, Cox4i2*) and SMC (*Acta2, Myh11, Tagln, Actg2*). **(B)** Violin plots showing the expression of *Ednra*, *Gpc3*, *Npy1r*, *Pln*, and *Atp1b2* in each cell type of mouse cavernous tissues. **(C)** AUC scores of *Rgs5, Lbh, Endra, Gpc3, Npylr, Pln*, and *Atplb2*. **(D)** ATP1B2 (red)/a-SMA (green) staining in MCPs and aorta SMC. Scale bars, 25 µm. **(E)** LBH (red)/a-SMA (green) and LBH (red)/CD31(green) staining in MCPs, aorta SMC. Scale bars, 100 µm. The nuclei were stained with DAPI (blue). Percentage of relative markers expressed cells. MCPs, mouse cavernous pericytes; SMC, smooth muscle cells.

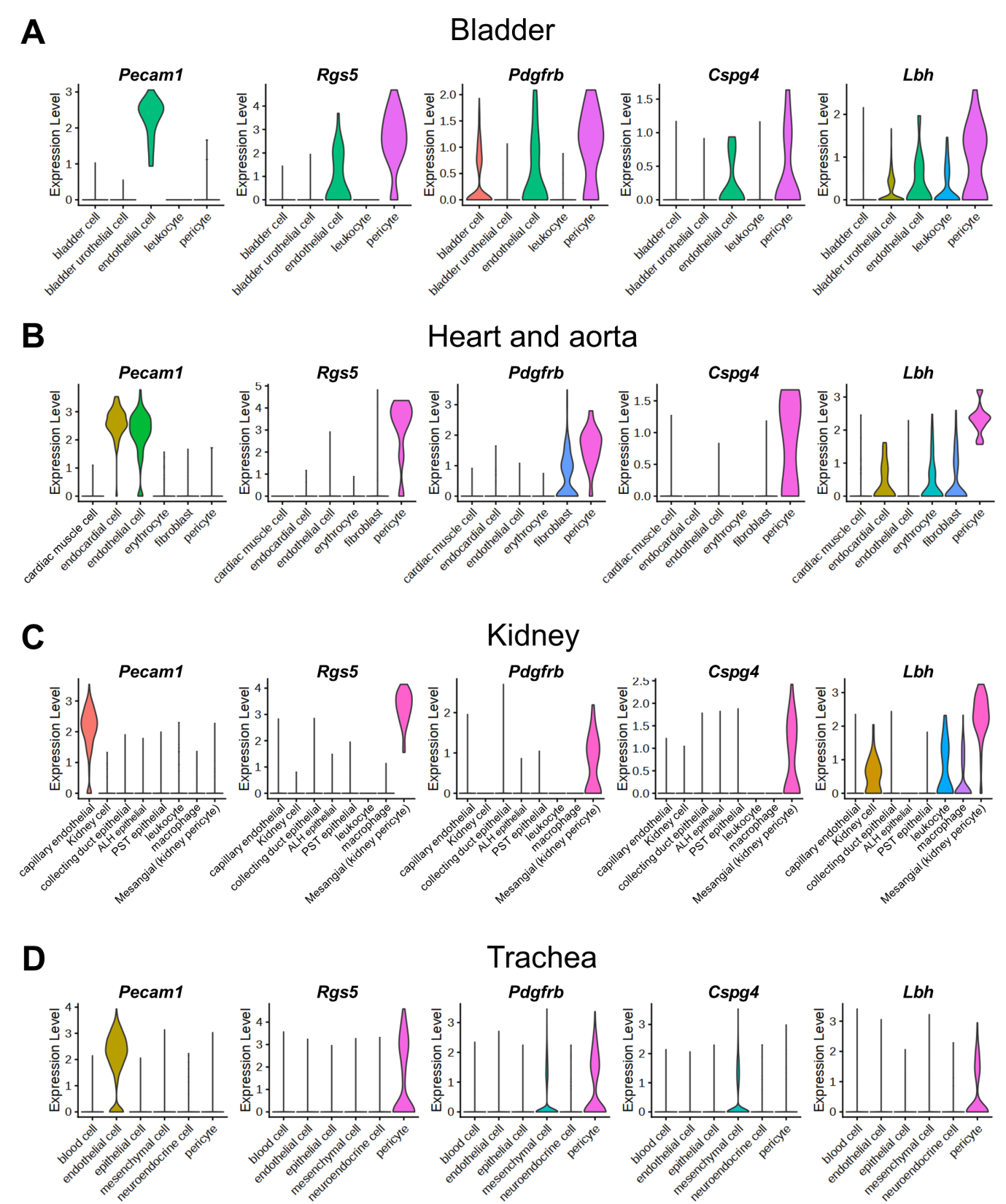

**Fig. S6. Expressions of *Lbh* in single-cell RNA sequencing data from tissues other than the penis.** ALH, Ascending limb of loop of Henle; PST, Proximal straight tubule. (**A-D**) Violin plots showing the expression of endothleial cell marker (*Pecam1*), known pericyte markers (*Rgs5, Pdgfrb, Cspg4*) and *Lbh* in mouse bladder (A), heart and aorta (**B**), kidney (**C**), trachea (**D**) from Tabula Muris.

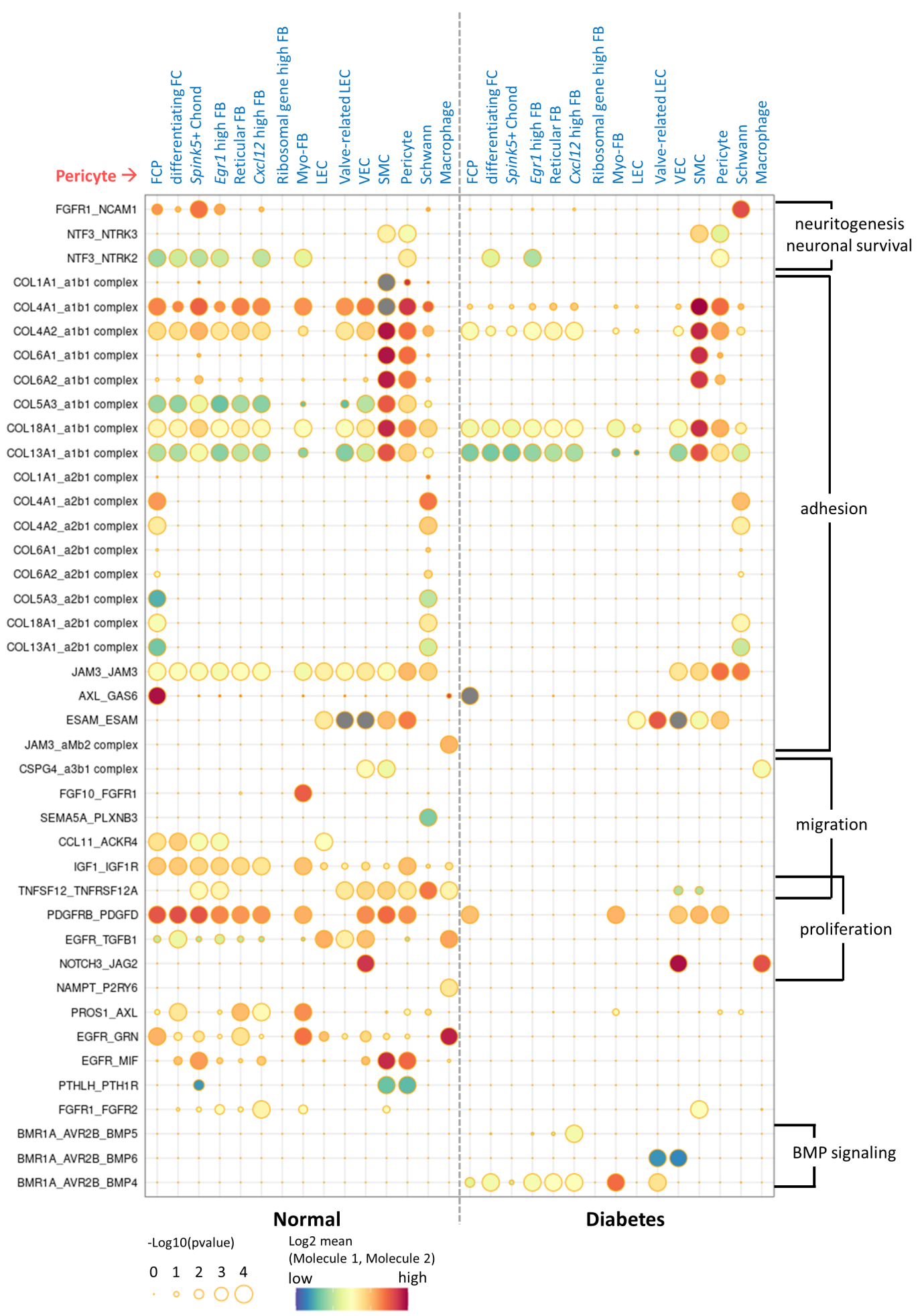

**Fig. S7. Cell-cell interactions between pericytes and other cell types showing significant differences in between normal and diabetes.** CellphoneDB dot plots showing ligand-receptor interactions between pericytes and other cell types in normal and diabetes. The plot shows all ligand-receptor interactions with significant differences between diabetes and normal. P-values are indicated as circle sizes. The means of the average expression level of the interaction are indicated by color.

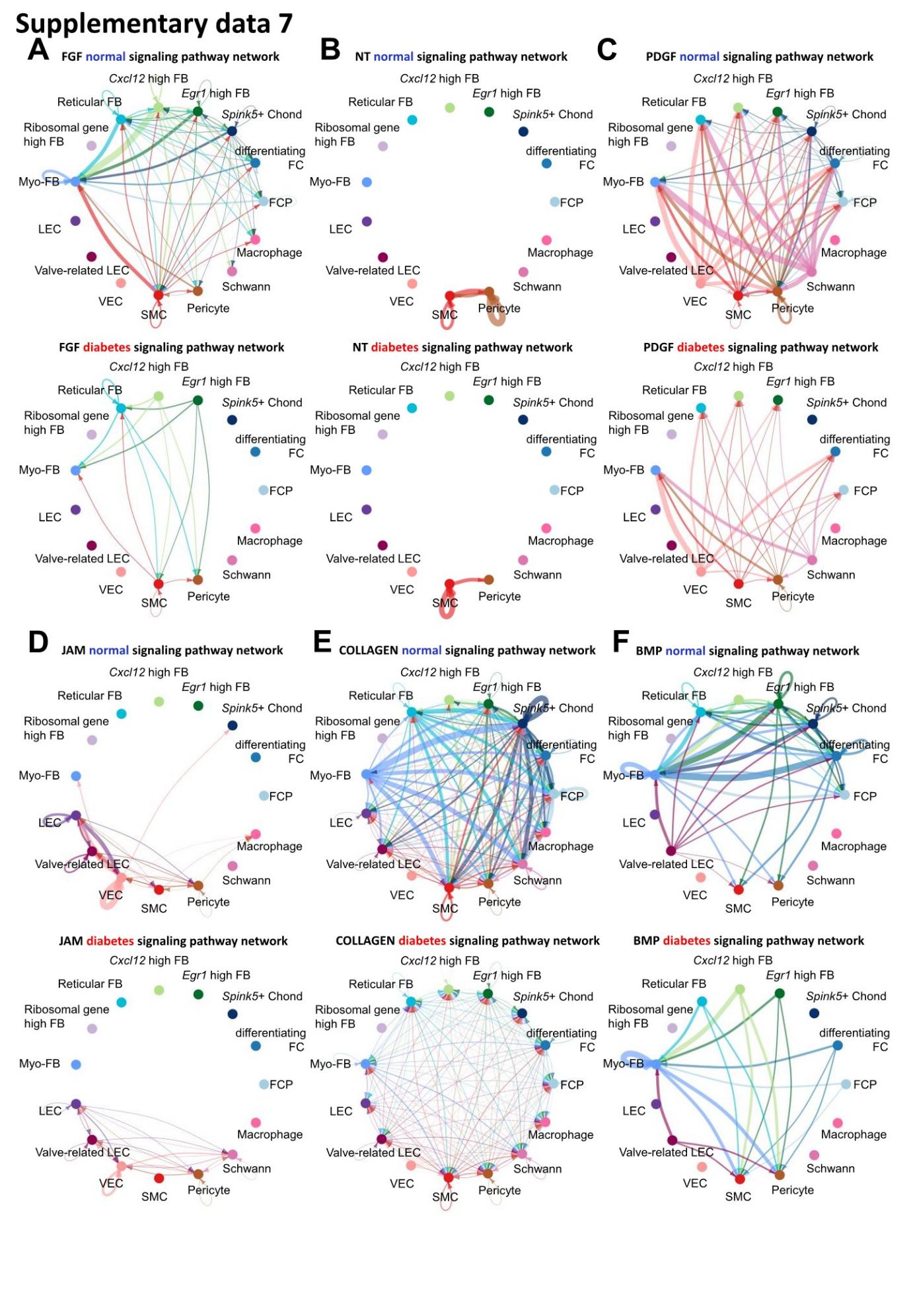

**Fig. S8. Cell-cell communication between cell types in normal and diabetes using CellChat.** (**A**) The inferred FGF signaling network in normal and diabetes. (**B**) The inferred NT signaling network in normal and diabetes. (**C**) The inferred PDGF signaling network in normal and diabetes. (**D**) The inferred JAM signaling network in normal and diabetes. (**E**) The inferred COLLAGEN signaling network in normal and diabetes. (**F**) The inferred BMP signaling network in normal and diabetes. The width of line represents the communication probability. The color of line matches the sender of the signal.

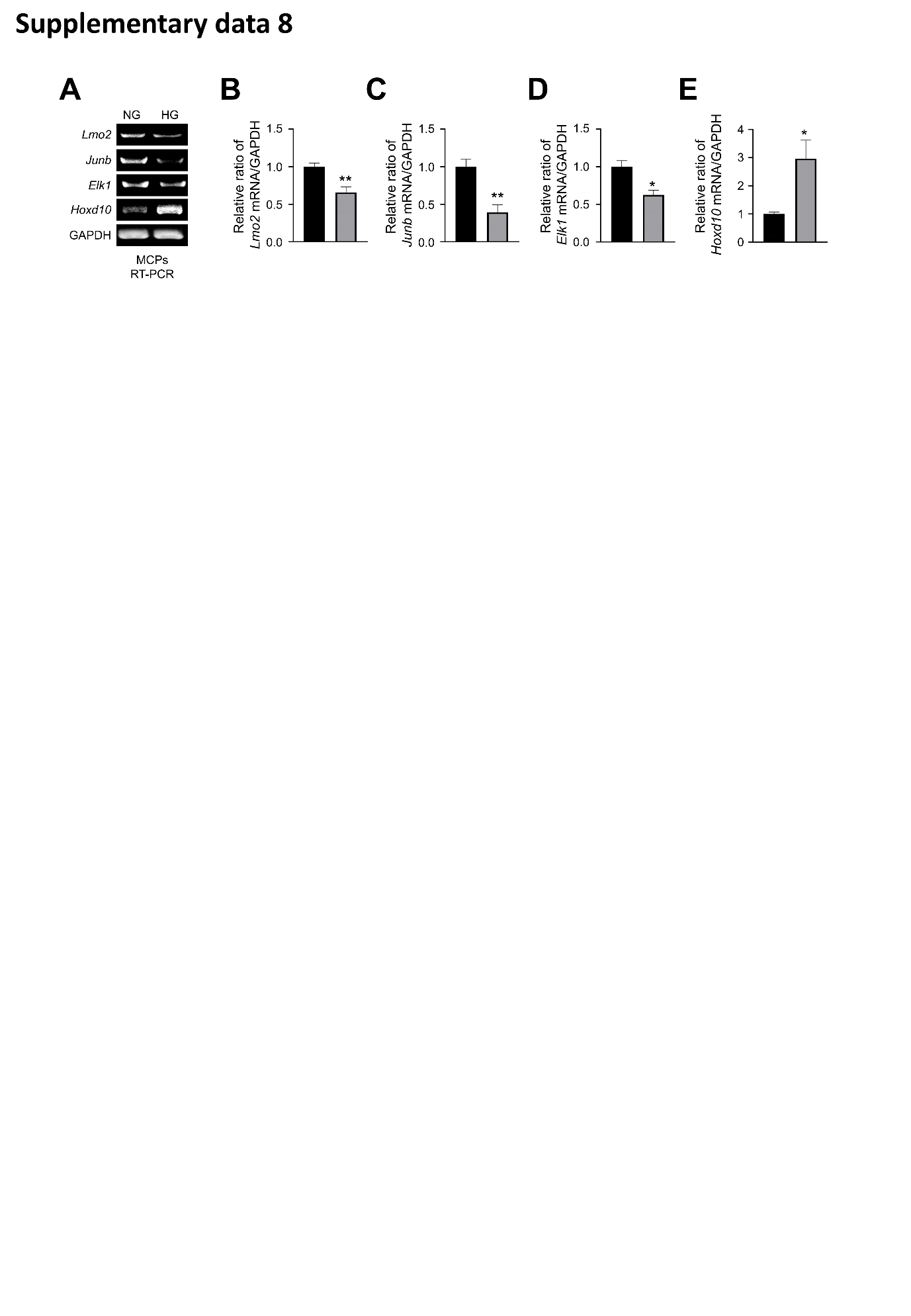

**Fig. S9. RT-PCR validation of differentially expressed genes from single cell RNA sequencing analyses in MCPs exposed to NG or HG conditions for 72 hours.** (**A**) Representative RT-PCR for *Lmo2, Junb, Elk1, Hoxd10* in MCPs. (**B-E**) Normalized band intensity values (n=4). *P < 0.05 vs. NG group. The relative ratio of the NG group was arbitrarily set to 1. NG, normal glucose; HG, high glucose.

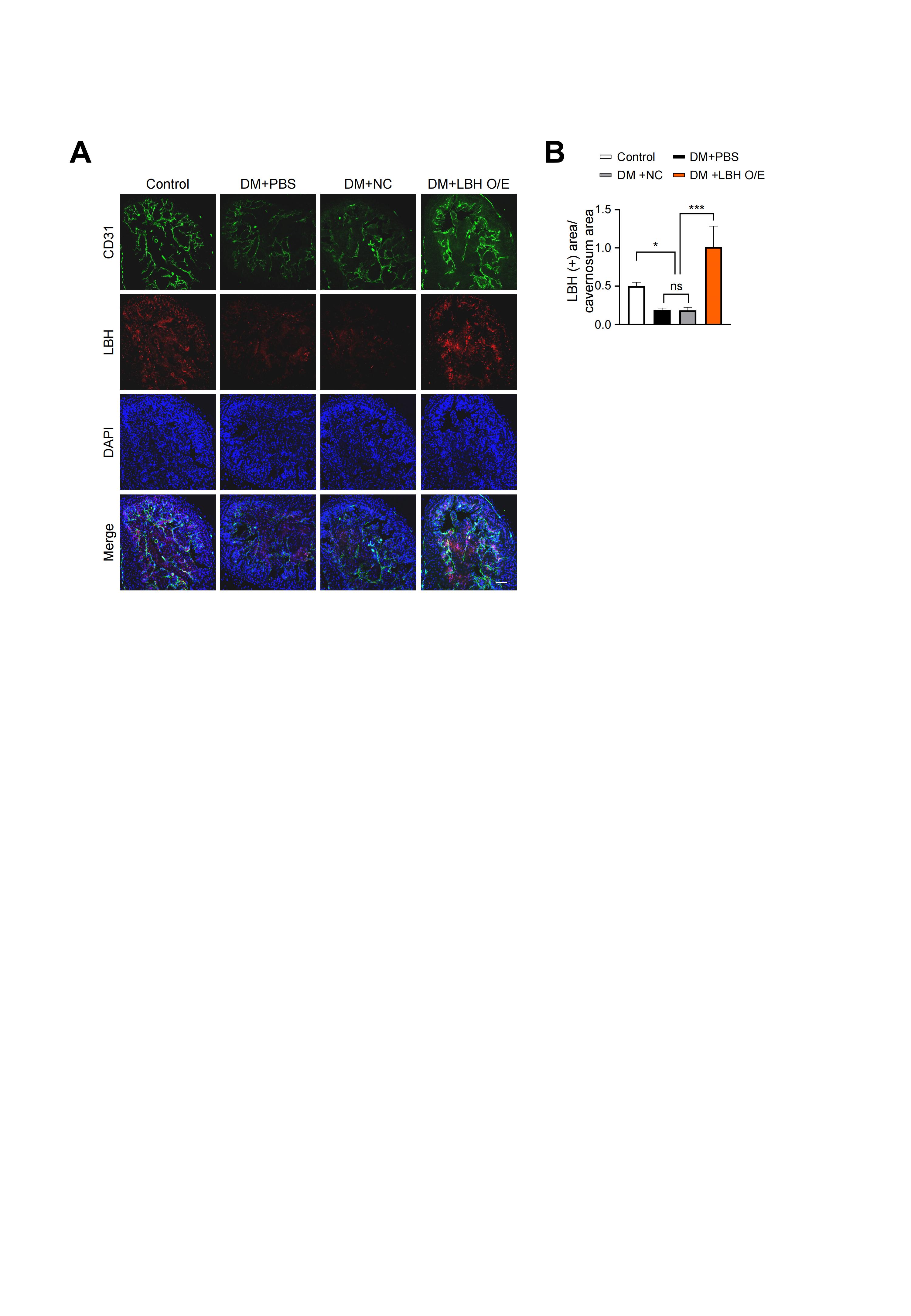

**Fig. S10. LBH immunofluorescence staining in corpus cavernosum tissues after infection with lentiviruses containing ORF mouse clone of *Lbh*.** (**A**) immunofluorescence staining of CD31 (green) and LBH (red) in corpus cavernosum tissues treated with lentiviruses ORF control particles (NC) and ORF clone of mouse *Lbh* (LBH O/E) in diabetic mice. Nuclei were labeled with DAPI (blue). Scale bars, 50 µm. (**B**) Quantification of LBH positive area by Image J, and results are presented as means ± SEM (n = 4). The relative ratio in the Control group was defined as 1. *P < 0.05; ***P < 0.001. DM, diabetes mellitus; DAPI, 4,6-diamidino-2-phenylindole.

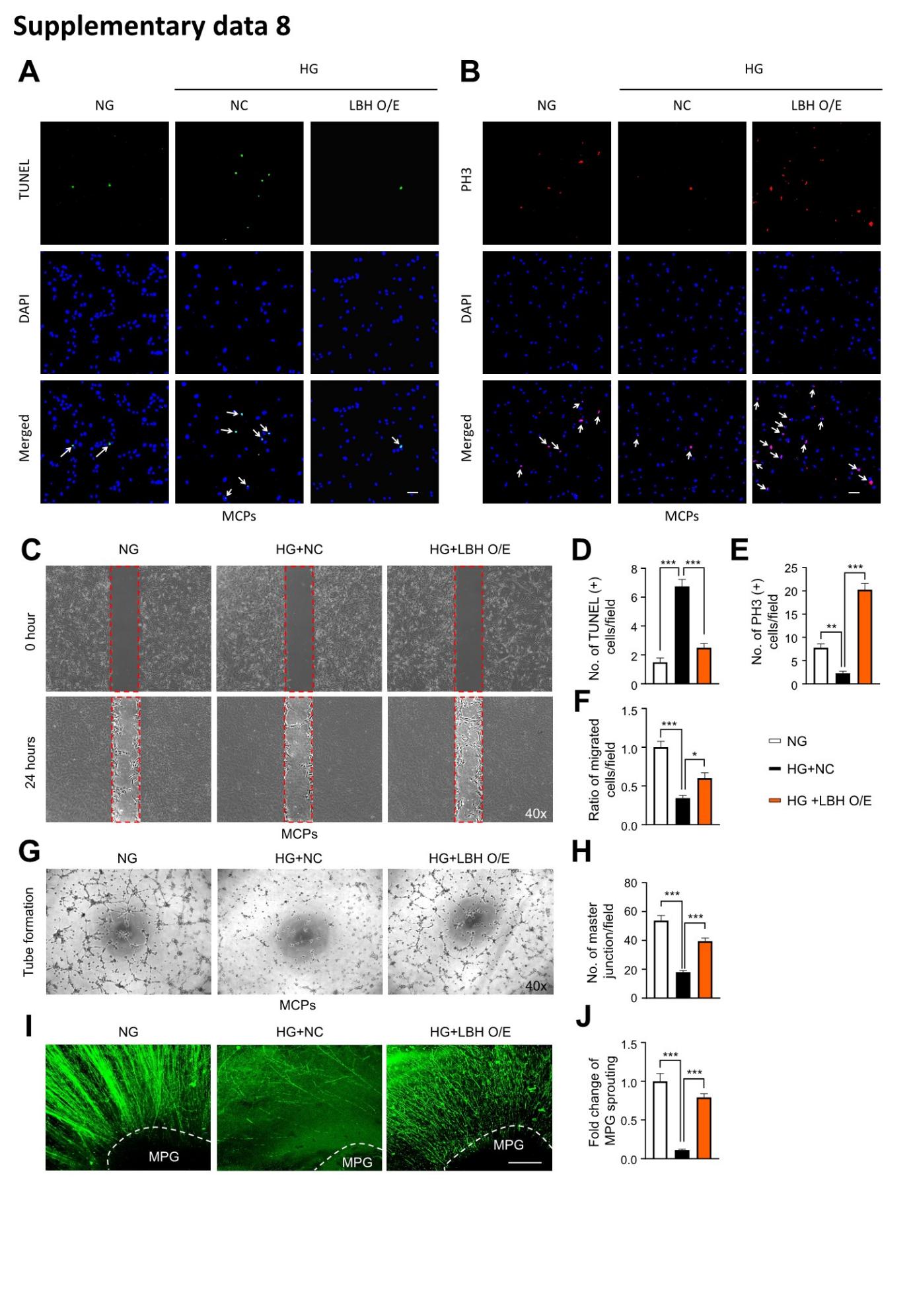

**Fig. S11. LBH enhances pericyte angiogenesis and MPG neurite sprouting under high glucose conditions.** (**A, B, C, G**, and **I**) TUNEL assay (**A**, green), immunofluorescence staining of PH3 (**B**, red), migration (**C**), tube formation (**G**) in MCPs, and immunofluorescence staining with neurofilament in MPG tissues (**I**, green) treated with lentiviruses ORF control particles (NC) and ORF clone of mouse *Lbh* (LBH O/E) under HG conditions. Nuclei were labeled with DAPI (blue). Scale bars, 50 µm. (**D, E, F, H**, and **J**) Quantification of number of TUNEL positive cells (arrow indicated, **D**), PH3 positive cells (arrow indicated, **E**), ratio of migrated cells (cells in red frame dot line, **F**), master junctions (**H**), and fold change of MPG sprouting (**J**) by Image J, and results are presented as means ± SEM (n = 4). The relative ratio in the NG group was defined as 1. *P < 0.05; **P < 0.01; ***P < 0.001. TUNEL, terminal deoxynucleotidyl transferase-mediated deoxyuridine triphosphate nick end labeling; NG, normal glucose; HG, high glucose; DM, diabetes mellitus; DAPI, 4,6-diamidino-2-phenylindole; MPG, major pelvic ganglion.

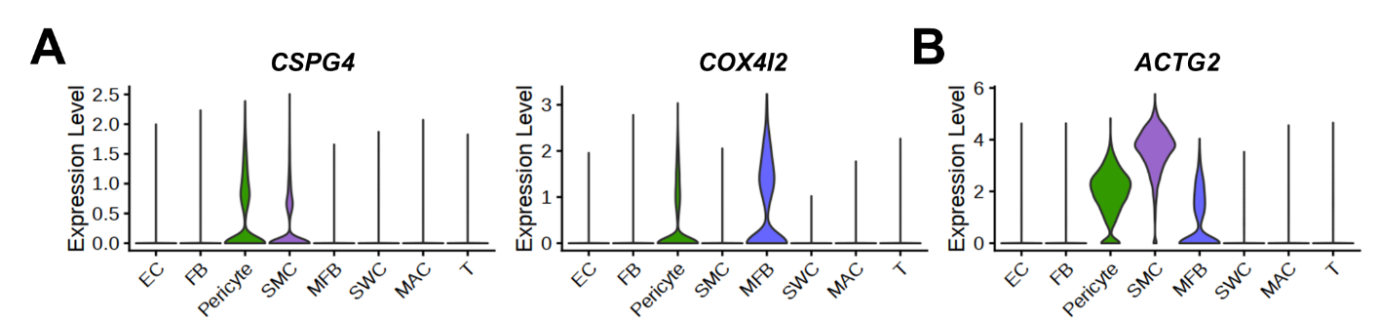

**Fig. S12. Expression of marker genes of pericyte and smooth muscle cells in human penis single cell RNA sequencing data. (A)** Violin plots showing known pericyte marker genes, *CSPG4* and *COX4I2*. **(B)** A violin plot showing known SMC marker genes, *ACTG2*. EC, endothelial cells;FB, Fibroblasts; SMC, Smooth muscle cells; MFB, Myofibroblast; SWC, Schwann cells; MAC, Macrophage;T, T cell.

**The mouse angiogenesis array coordinates are shown in Table S1**

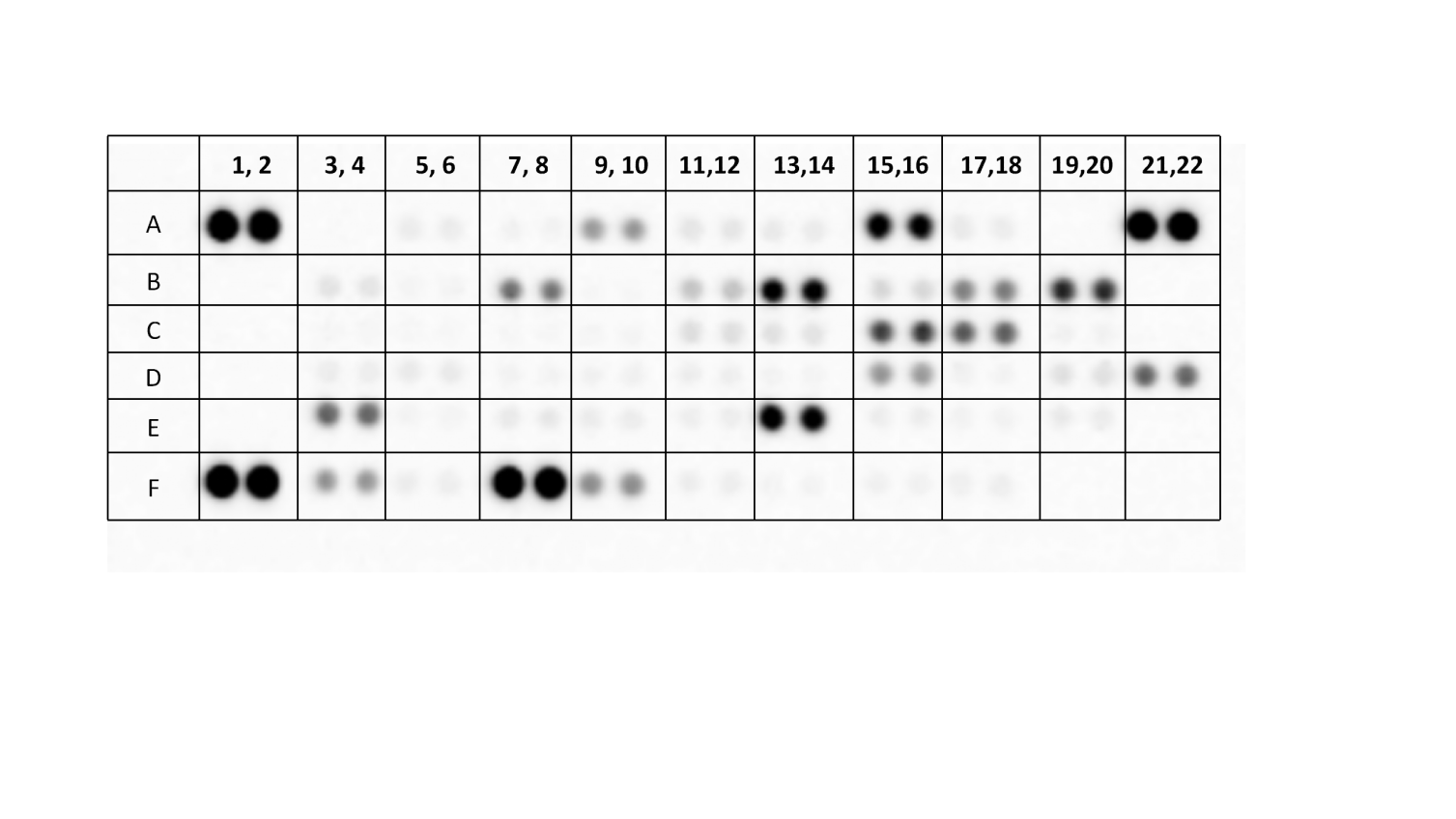

| **Table S1.** Relative density of mouse angiogenesis array spots | | | | |
| --- | --- | --- | --- | --- |
| **Coordinate** | **Analyte/Control** | **Density in Normal** | **Density in DM** | **Ratio (normal/DM)** |
| A1, A2 | Reference Spots | 40198 | 39327 | 0.98 |
| A5, A6 | ADAMTS1 | 1355 | 1155 | 0.85 |
| A7, A8 | Amphiregulin | 410 | 543 | 1.32 |
| **A9, A10** | **Angiogenin** | **5071** | **6522** | **1.29** |
| A11, A12 | Angiopoietin-1 | 1592 | 1561 | 0.98 |
| A13, A14 | Angiopoietin-3 | 1283 | 1153 | 0.90 |
| A15, A16 | Coagulation Factor III | 25262 | 23574 | 0.93 |
| A17, A18 | CXCL16 | 1090 | 1174 | 1.08 |
| A21, A22 | Reference Spots | 42612 | 40543 | 0.95 |
| B3, B4 | Cyr61 | 1660 | 1672 | 1.01 |
| B5, B6 | DLL4 | 427 | 524 | 1.23 |
| B7, B8 | DPPIV | 7955 | 9405 | 1.18 |
| B9, B10 | EGF | 90 | 110 | 1.22 |
| **B11, B12** | **Endoglin (CD105)** | **5743** | **3330** | **0.58** |
| B13, B14 | Endostatin/Collagen XVIII | 20887 | 21221 | 1.02 |
| B15, B16 | Endothelin-1 | 1765 | 1638 | 0.93 |
| B17, B18 | FGF acidic | 9771 | 9275 | 0.95 |
| B19, B20 | FGF basic | 15057 | 16760 | 1.11 |
| C3, C4 | KGF | 118 | 170 | 1.44 |
| C5, C6 | Fractalkine | 150 | 131 | 0.87 |
| C7, C8 | GM-CSF | 244 | 300 | 1.23 |
| C9, C10 | HB-EGF | 363 | 390 | 1.07 |
| C11, C12 | HGF | 399 | 360 | 0.90 |
| C13, C14 | IGFBP-1 | 169 | 185 | 1.09 |
| **C15, C16** | **IGFBP-2** | **16754** | **13118** | **0.78** |
| C17, C18 | IGFBP-3 | 11454 | 12038 | 1.05 |
| C19, C20 | IL-1α | 97 | 94 | 0.97 |
| C21, C22 | IL-1β | 35 | 17 | 0.49 |
| D3, D4 | IL-10 | 1058 | 1150 | 1.09 |
| D5, D6 | IP-10 | 856 | 845 | 0.99 |
| D7, D8 | KC | 542 | 540 | 1.00 |
| D9, D10 | Leptin | 989 | 823 | 0.83 |
| D11, D12 | MCP-1 | 857 | 871 | 1.02 |
| D13, D14 | MIP-1α | 695 | 612 | 0.88 |
| **D15, D16** | **MMP-3 (pro and mature form)** | **1656** | **7314** | **4.42** |
| D17, D18 | MMP-8 (pro form) | 326 | 338 | 1.04 |
| D19, D20 | MMP-9 (pro and active form) | 1770 | 1999 | 1.13 |
| **Table S1.** Relative density of mouse angiogenesis array spots (Continued) | | | | |
| **D21, D22** | **NOV (IGFBP-9)** | **14787** | **10986** | **0.74** |
| **E3, E4** | **Osteopontin (OPN)** | **15851** | **9510** | **0.60** |
| E5, E6 | PD-ECGF | 304 | 388 | 1.28 |
| E7, E8 | PDGF-AA | 1008 | 1226 | 1.22 |
| E9, E10 | PDGF-AB/PDGF-BB | 817 | 1055 | 1.29 |
| E11, E12 | Pentraxin-3 | 852 | 1065 | 1.25 |
| **E13, E14** | **Platelet Factor 4** | **19577** | **23907** | **1.22** |
| E15, E16 | PlGF-2 | 625 | 687 | 1.10 |
| E17, E18 | Prolactin | 476 | 550 | 1.16 |
| E19, E20 | Proliferin | 604 | 831 | 1.38 |
| F1, F2 | Reference Spots | 40870 | 41967 | 1.03 |
| F3, F4 | SDF-1 | 7034 | 7236 | 1.03 |
| F5, F6 | SerpinE1 | 1395 | 1096 | 0.79 |
| F7, F8 | Serpin F1 | 36552 | 38994 | 1.07 |
| **F9, F10** | **Thrombospondin-2 (TSP-2)** | **13897** | **7254** | **0.52** |
| F11, F12 | TIMP-1 | 474 | 884 | 1.86 |
| F13, F14 | TIMP-4 | 158 | 413 | 2.61 |
| F15, F16 | VEGF | 292 | 666 | 2.28 |
| F17, F18 | VEGF-B | 517 | 832 | 1.61 |
| F19, F20 | Negative Control | 15 | 17 | 1.13 |

| **Table S2.** Physiologic and metabolic parameters: 2 weeks after treatment with PBS, NC, LBH O/E | | | | |
| --- | --- | --- | --- | --- |
|  | | **STZ-induced diabetic mice (DM)** | | |
|  | **Control** | **PBS** | **NC** | **LBH O/E** |
| **Body weight (g)** | 31.7±1.1 | 24.1±1.9* | 24.9±0.7* | 25.3±0.8* |
| **Fasting glucose (mg/dl)** | 102.6±4.1 | 549.2±41.7* | 559.2±39.1* | 548.2±39.9* |
| **Postprandial glucose (mg/dl)** | 171.2±37.9 | 588±13.2* | 591.2±12.1* | 587±13.7* |
| **MSBP (mm Hg)** | 100.7±5.6 | 105.8±2.8 | 100.0±4.4 | 104.6±4.2 |
| Values are the mean ± SEMs for n=5 animals for each group. STZ, streptozotocin; MSBP, Mean systolic blood pressure; NC, lentiviruses ORF control particles; LBH O/E, ORF clone of mouse Lbh lentiviruses, **P* < 0.05 vs. Control group | | | | |
